# Binding of Monomeric and Polymeric Alzheimer’s Aβ peptides to Exosomes

**DOI:** 10.1101/2021.08.06.455470

**Authors:** Christina Coughlan, Jared Lindenberger, Jeffrey Jacot, Noah Johnson, Paige Anton, Shaun Bevers, Michael Graner, Huntington Potter

## Abstract

Exosomes are secreted by every cell in our body under both physiological and pathological conditions. They travel in the blood, CSF, and all studied biofluids. Their biological roles have been reported to include delivery of important physiological cargo between organs and cells, clearance of toxic proteins; maintenance of cellular stasis, and the propagation of disease pathology. In the case of Alzheimer’s disease (AD) exosomes have been shown to carry pathological proteins such as amyloid, yet the specificity of this association of amyloid and exosomes is unclear. To address this deficiency, we utilized Isothermal Titration Calorimetry (ITC) to measure the binding of amyloid to exosomes. Here we report that Aβ40 and Aβ42 bind to exosomes in a saturable and endothermic manner, a phenomenon not observed with the scrambled versions of either peptide. This points to this interaction being more specific than previously understood, and to amyloid associated with exosomes as an important pool of this peptide in the plasma.

## Introduction

Exosomes are 50-150 nm diameter vesicles that are secreted by every cell in the body. The generation of exosomes starts in the endosomal system with early endosomes maturing into late endosomes or forming multivesicular bodies (MVB’s) [1–4]. MVB’s contain within them Intraluminal Vesicles (ILV’s) formed by endosomal membrane invagination into the lumen of the endosome, a process modulated by ESCRT complexes, Rab GTPases, as well as ESCRT independent pathways [5–7]. These ILV’s can be targeted to the lysosome for degradation, or secreted from the cell as exosomes, a process that occurs when the MVB’s fuse with the plasma membrane under both normal and pathological conditions. When secreted as exosomes they can travel in biofluids and are thus capable of having effects both remotely and locally.

To date exosomes have been reported to carry 41,860 proteins, 1,116 lipids and 7,540 RNA molecules [8] with the cargo and composition of exosomes thought to reflect the physiological and pathological status of the parent cell [6]. These numbers represent the total molecules identified and submitted to the various databases. Any individual exosome of ~ 100nm diameter likely carries approximately 100 proteins on the interior and perhaps 100 proteins on the exterior surfaces [8] [9]. Once secreted exosomes can enter cells by selectively binding to surface components, some of which have been identified. They can also enter cells by fusing with lipid bilayers. Both routes of entry allow exosomes to share their cargo locally and remotely, pointing to their potential role in both physiological and pathological processes. Thus, deciphering cargo that is selectively loaded into and onto exosomes under normal physiological and pathological conditions is imperative.

While exosomes have been reported to carry Amyloid, there is little in the way of knowledge as to whether this Amyloid is inside or outside the exosome or how specific this loading of Amyloid cargo is for exosomes. In this work, using Isothermal Titration Calorimetry (ITC), we measured the specificity with which amyloid is attached to intact exosomes in an attempt to decipher if Amyloid transport is a more selective process than previously envisioned. We found that Aβ40 and Aβ42 bound in an endothermic and saturable manner to plasma exosomes, a phenomenon not observed for the scrambled versions of either peptide. In addition Atomic Force Microscopy (AFM) showed that amyloid has a fibrillar form when exosomes and Aβ42 are incubated together. This points to Amyloid/Aβ having a more selective interaction with exosomes than previously considered and highlights the importance of measuring this pool of amyloid in the plasma.

## Materials and Methods

### Peptides

Recombinant human Aβ40 NaOH salt and Aβ42 NaOH salt were purchased from rPeptide. Aβ peptides were reconstituted in DPBS, pH 7.4 at a 200 μM concentration, sonicated for 10 minutes, flash-frozen in liquid nitrogen, and stored at −80C until further use.

### Binding of Aβ42 to exosomes

Eight preparations of exosomes were generated from 250μls of plasma each by precipitation using EQULTRA-20A-1 precipitation reagent (Systems Biosciences) as described previously [10]. Each resultant exosomal pellet was resuspended in 200μls of H_2_0 to which 10μls of each of the following was added: PBS, 0.2μM, 2μM or 20μM Aβ42 in monomeric or fibrillar form for final concentrations of 0 (PBS only), 0.01 μM, 0.1 μM and 1 μM. This mixture was incubated with rotation overnight at 37°C. The exosomes were then reprecipitated from this solution and the exosomal pellets lysed in 100 μls of 0.05M Glycine and centrifuged @4,000g for 10mins. To the supernatant of this centrifugation 15 μls of 1M Tris pH8.0, 25 μls of 3%BSA/PBS and 180 μls mPERS were added along with Protease Phosphatase inhibitors. This exosomal lysate was then frozen (−80°C) and thawed (37°C) ×2 and analyzed using the 3 Plex assay and SIMOA^®^ platform (Quanterix) as outlined by the manufacturers protocols.

### Isothermal Titration Calorimetry (ITC) Procedure

Isothermal titration calorimetry (ITC) experiments were performed at 25 °C using a Microcal ITC200 MicroCalorimeter (Malvern Panalytical). For the ITC experiment, the cell contained the peptides at 20 μM in PBS buffer and the syringe contained 2.31×10^11^ particles/ml of exosomes in PBS buffer. The volume of the first injection was 0.4 μl over 0.8 s (this initial injection was excluded from data analysis). The following injections were 2.0 μl over 4 seconds (19 total) with an injection spacing of 180 seconds. The reference power was set to 10 μcal/second in high feedback mode. The syringe stirring speed was set to 750 rpm. Data analyzed using Origin 7.0 software (OriginLab).

### Sample preparation of the Aβ42/exosomes for Atomic Force Microscopy (AFM)

Exosomes were generated by precipitation using EQULTRA-20A-1 (Systems Biosciences) and methods described previously [10]. Each resultant exosomal pellet was resuspended in 200μls of PBS to which 10μls of each of the following was added: PBS or 20μM Aβ42. This mixture was incubated with rotation overnight at 37°C. The exosomes were then reprecipitated from this solution and the pelleted versus supernatant material was analyzed using Atomic Force Microscopy (AFM).

### Atomic Force Microscopy (AFM)

AFM images were acquired with a JPK NanoWizard 4a AFM (JPK Instruments USA, Carpinteria, CA) with silicon-nitride cantilevers with a triangular tip and a nominal spring constant of 0.07 N/m (MLCT, Bruker AFM Probes, Camarillo, CA) run in either contact mode or AC (tapping) mode. For sample preparation, 50 μl of sample was pipetted onto a clean, freshly cleaved mica surface and dried at 50°C for 1 hour.

## Results

Amyloid can be measured, associated with exosomes, when isolated from plasma [1, 11–13]. In order to determine if Aβ binds to exosomes in a more specific manner we incubated exosomes with Aβ42 at a range of concentrations and used the SIMOA 3 Plex assay (Quanterix) to quantify the amount of Aβ42 bound. Of interest was that incubating exosomes with 0.1μM monomeric Aβ42 revealed 4x the amount of monomeric Aβ42 being attached to the exosomes compared to when fibrillar Aβ42 was utilized (1B). At 1μM Aβ42 concentration, monomeric Aβ42 was above the level of detection of the assay, whereas fibrillar Aβ42 binding was still measurable (1A).

To determine if monomeric Aβ40 and Aβ42 bind directly to exosomes, we utilized isothermal titration calorimetry (ITC) in which exosomes were titrated into a solution containing the Aβ peptides. Titration of exosomes into the Aβ42 peptide yielded an endothermic isotherm profile showing clear saturation of binding (Figure 2A). Titration of exosomes into the Aβ40 also yielded an endothermic isotherm profile (Figure 2B), similar to Aβ42, however, saturation of binding was never reached (Figure 2B). This may be due in part to the shorter peptide length having a lower affinity for the exosomes, thus increasing the amounts of exosomes needed to reach saturation. In contrast, both scrambled versions of the peptides exhibit exothermic isotherm patterns (Figure 2C and 2D). These patterns very closely match the control experiment of titrating exosomes into PBS buffer (Figure 2E). This indicates that the scrambled versions of both the monomeric Aβ40 and Aβ42 are unable to interact with the exosomes directly.

**Fig 1:**
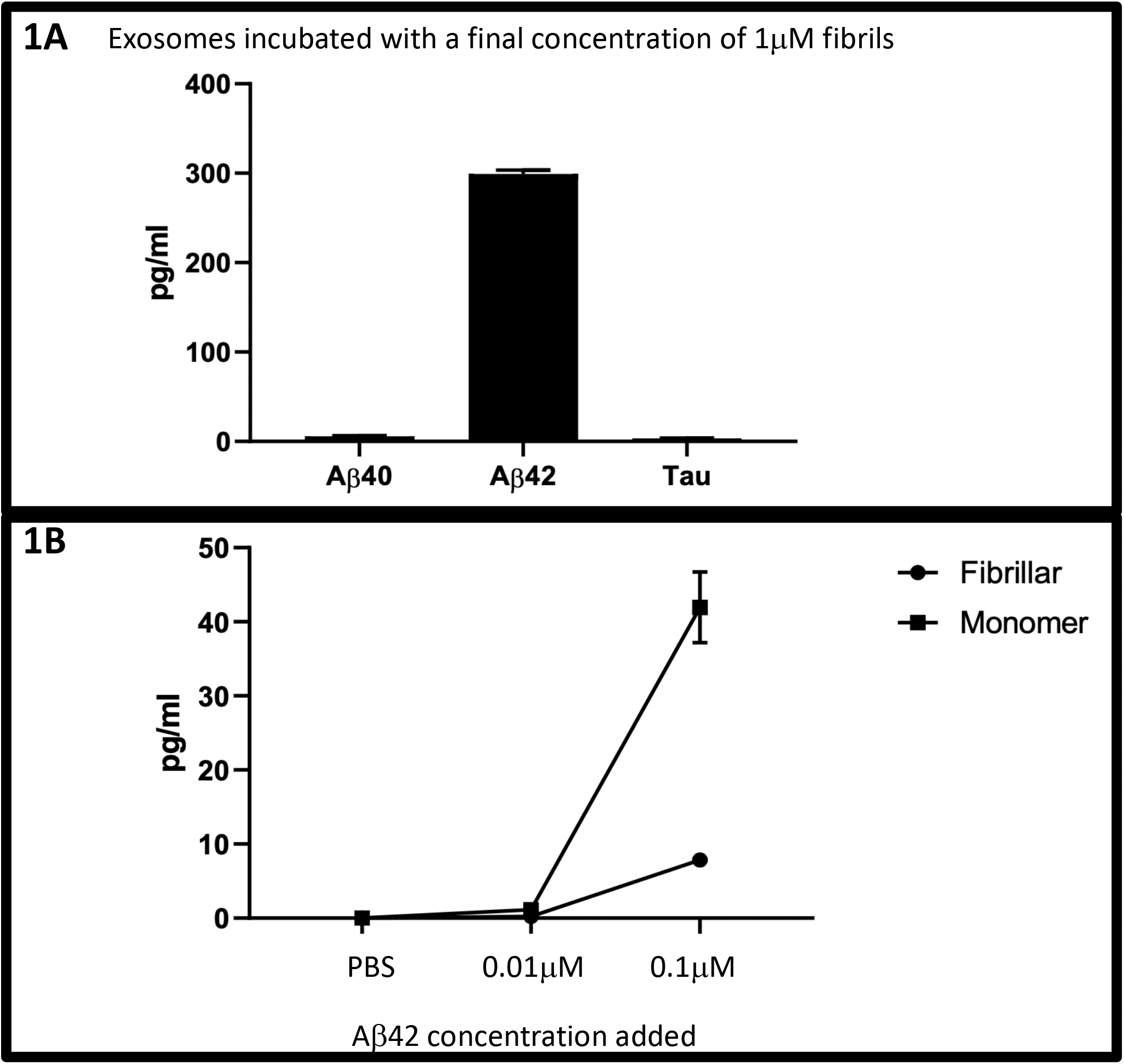
Binding of Aβ42 to exosomes: To determine if Aβ42 binds to exosomes we incubated Aβ42 at various concentrations with exosomes and used the SIMOA 3 Plex assay (Quanterix) to measure the amount of Aβ42 detectably bound. Of interest is that at an incubation concentration of 0.1μM monomeric Aβ42, exosomes had 4x the amount of Aβ42 associated with them compared to when the same concentration of fibrillar Aβ42 (Figure 1B) was utilized. Adding a concentration of 1μM Aβ42, bound monomeric Aβ42 was above the level of detection of the assay, whereas fibrillar Aβ42 binding was still measurable (Figure 1A). Aβ40 and Tau were not detectable, which was expected given only Aβ42 was incubated with the exosomes.

**Fig 2:**
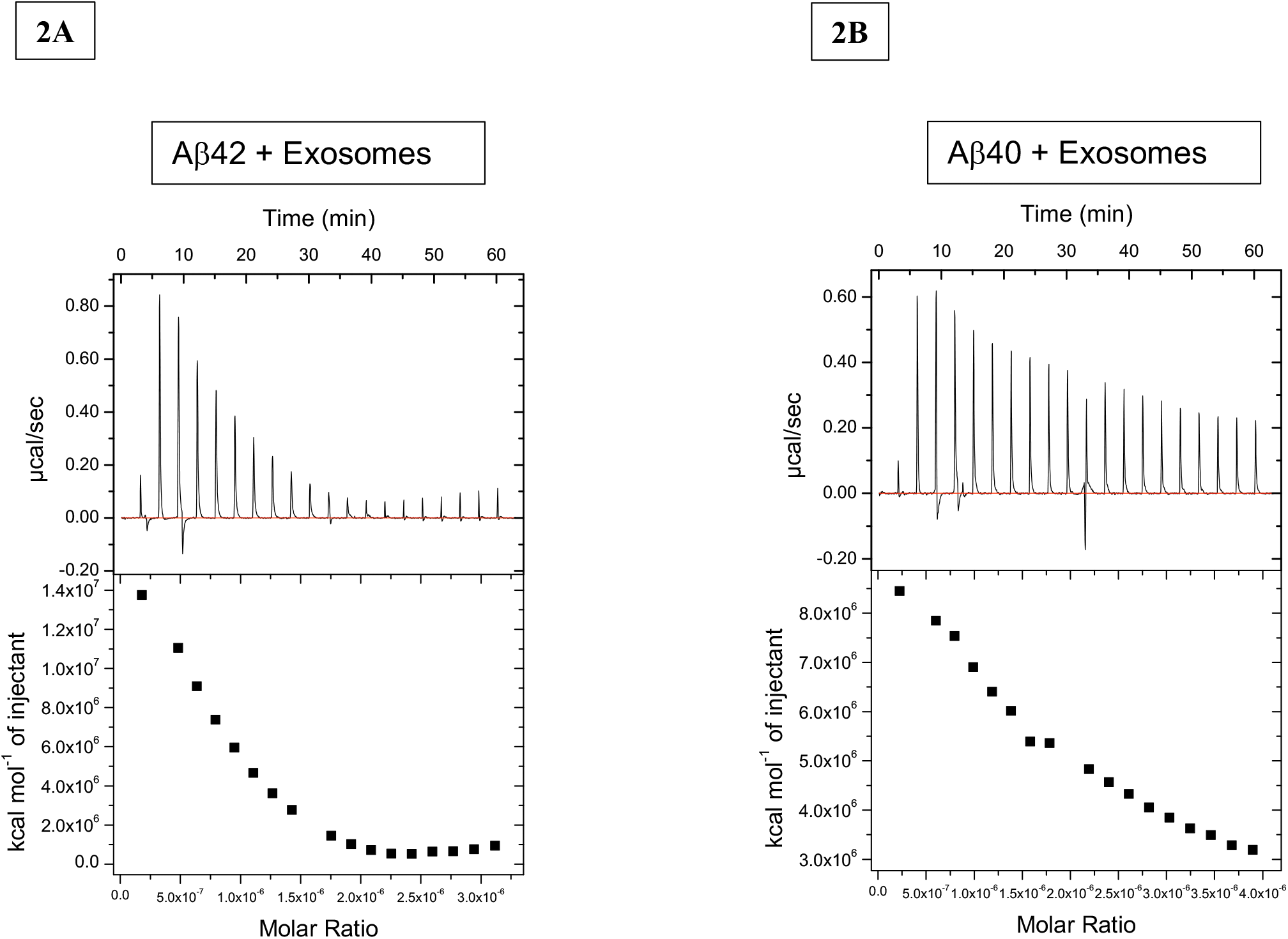
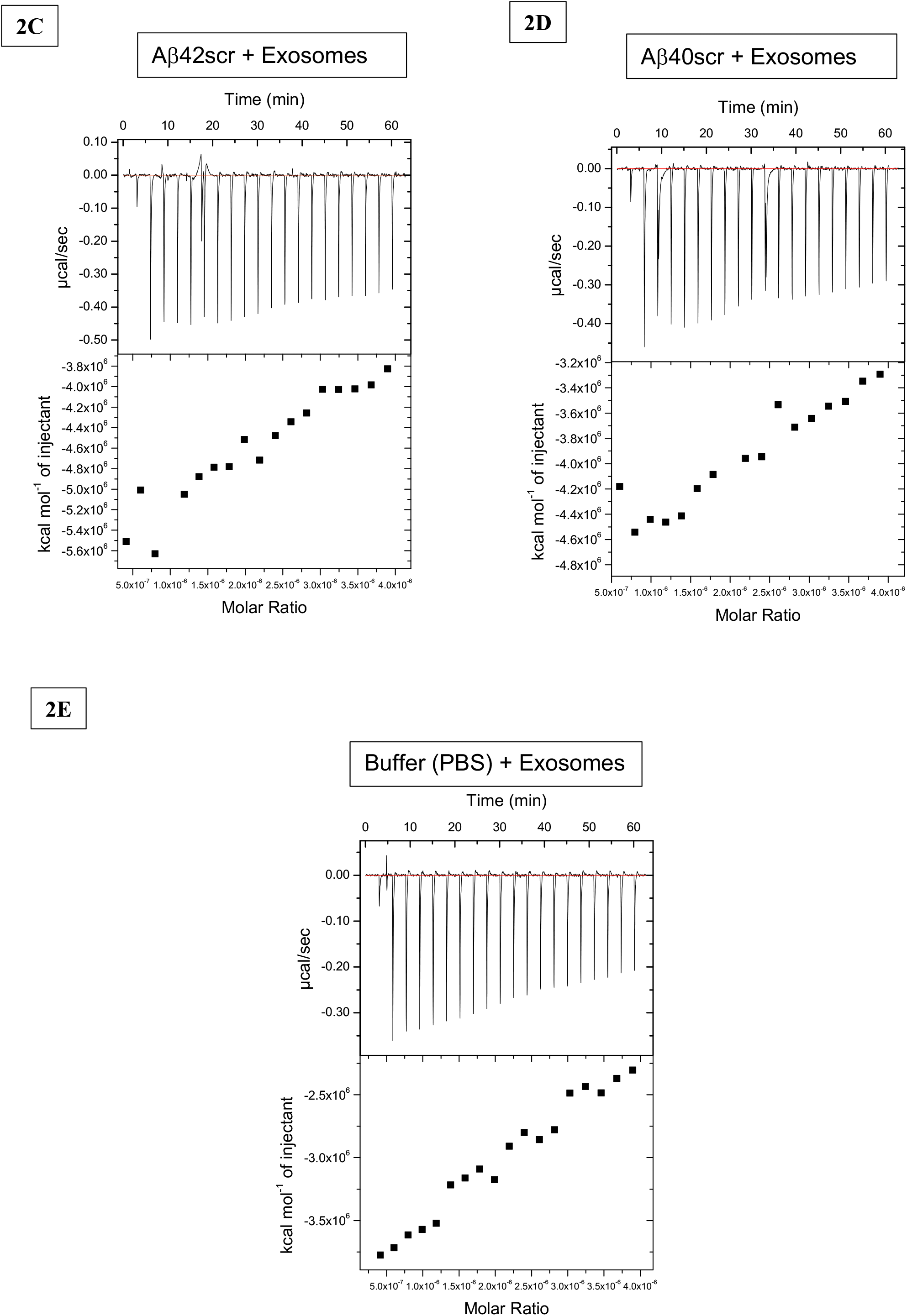
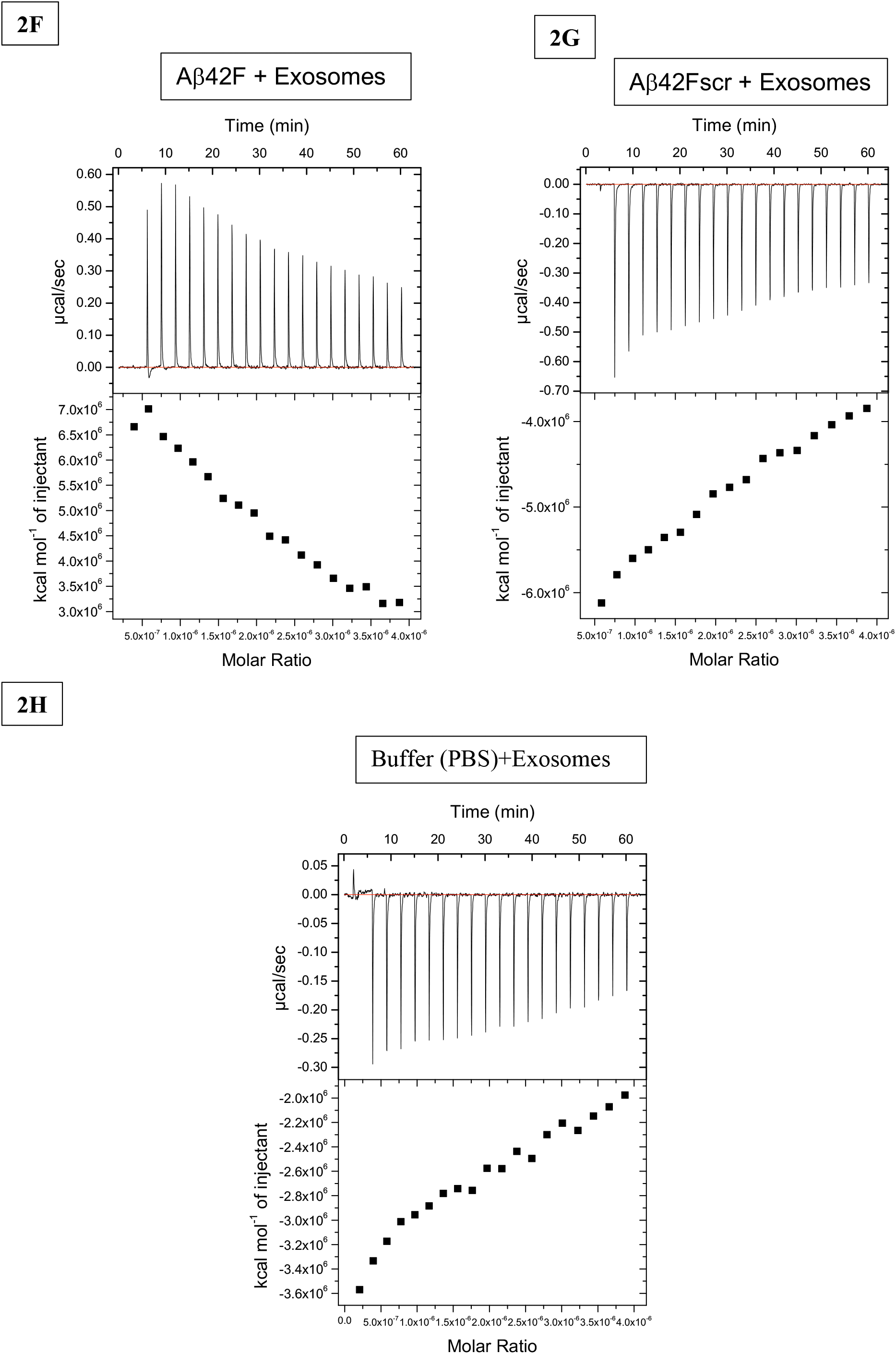
Assessment of Amyloid binding to exosomes using Isothermal Titration Calorimetry (ITC). To determine if monomeric Aβ40 and Aβ42 bind directly to exosomes, we utilized isothermal titration calorimetry. Exosomes were titrated into a solution containing the various types and forms of Aβ peptides Aβ42 (Figure 2A), Aβ40 (Figure 2B), Aβ42scr (Figure 2C), Aβ40scr (Figure 2D), Buffer alone (Figure 2E), Aβ42F (Figure 2F), Aβ42Fscr (Figure 2G) and Buffer alone for the F assay (Figure 2H). The exosomes used in the titration are heterogeneous, meaning all have different diameters and surface areas. Therefore, the data cannot be fit to determine exact thermodynamic binding parameters given there is likely quite a variety of stoichiometries with the number of different exosome species in the sample. Therefore, we can only approximate a Molar ratio of Amyloid (20 μM) to exosomes (2.31×10^11 particles/ml) which helped us assess that the peptides exhibited one pattern of binding, while the scrambled versions exhibited another pattern of binding similar to the control reaction of exosome titration into buffer. Taken together, these data suggest that the exosomes are capable of binding the Aβ peptides in a direct and specific manner. Beacause the molar concentration of particles is small compared to the peptide this explains why the ratio is so small.

Additionally, fibrillar Aβ42 (Aβ42F) was tested to ascertain whether the peptide could still bind the exosomes in its fibrillar form. From the data we see that the Aβ42F can indeed bind to exosomes (Figure 2F) and exhibits the same endothermic binding profile as monomeric Aβ42. When the titration is performed with scrambled Aβ42Fscr, that was prepared in the same manner as the Aβ42F, we see an exothermic profile (Figure 2G), which is similar to the control titration with exosomes into PBS buffer (Figure 2H). This again suggest that only Aβ42, in this case Aβ42F, can bind to the exosomes, while the scrambled version cannot.

The exosomes used in the titration are heterogeneous, meaning all have different diameters and surface areas, and thus, different capacities to bind the peptide. Therefore, the data cannot be fit to determine exact thermodynamic binding parameters as we possibly have a variety of stoichiometries for the different exosome species in the sample. Therefore, we can conclude that the peptides exhibit one pattern of binding, while the scrambled versions exhibit another pattern of binding that is similar to a control reaction of exosome titration into buffer. Taken together, this data suggests that the exosomes are capable of binding the Aβ peptides in a direct and specific manner.

To visualize the exosome/Aβ42 interaction, exosomes and Aβ42 were incubated overnight, precipitated from that solution and imaged utilizing Atomic Force Microscopy (AFM) with AC mode at a frequency of 375Hz and amplitude around 900nm. Imaging showed that when Aβ42 and exosomes were mixed, visible aligned fibers formed with heights of 2-3 nm (Figure 3A). However, in the supernatant from the precipitation of the incubated exosome/Aβ42 sample, no fibers and only small, more monomeric structures of Aβ, were observed (Figure 3B). Precipitation of exosomes without Aβ42 present shows soft structures but no fibers (Figure 3C) while analysis of samples that resulted from precipitation of Aβ42 alone (no exosomes present) resulted in empty surfaces with only occasional small structures (Figure 3D).

**Figure 3:**
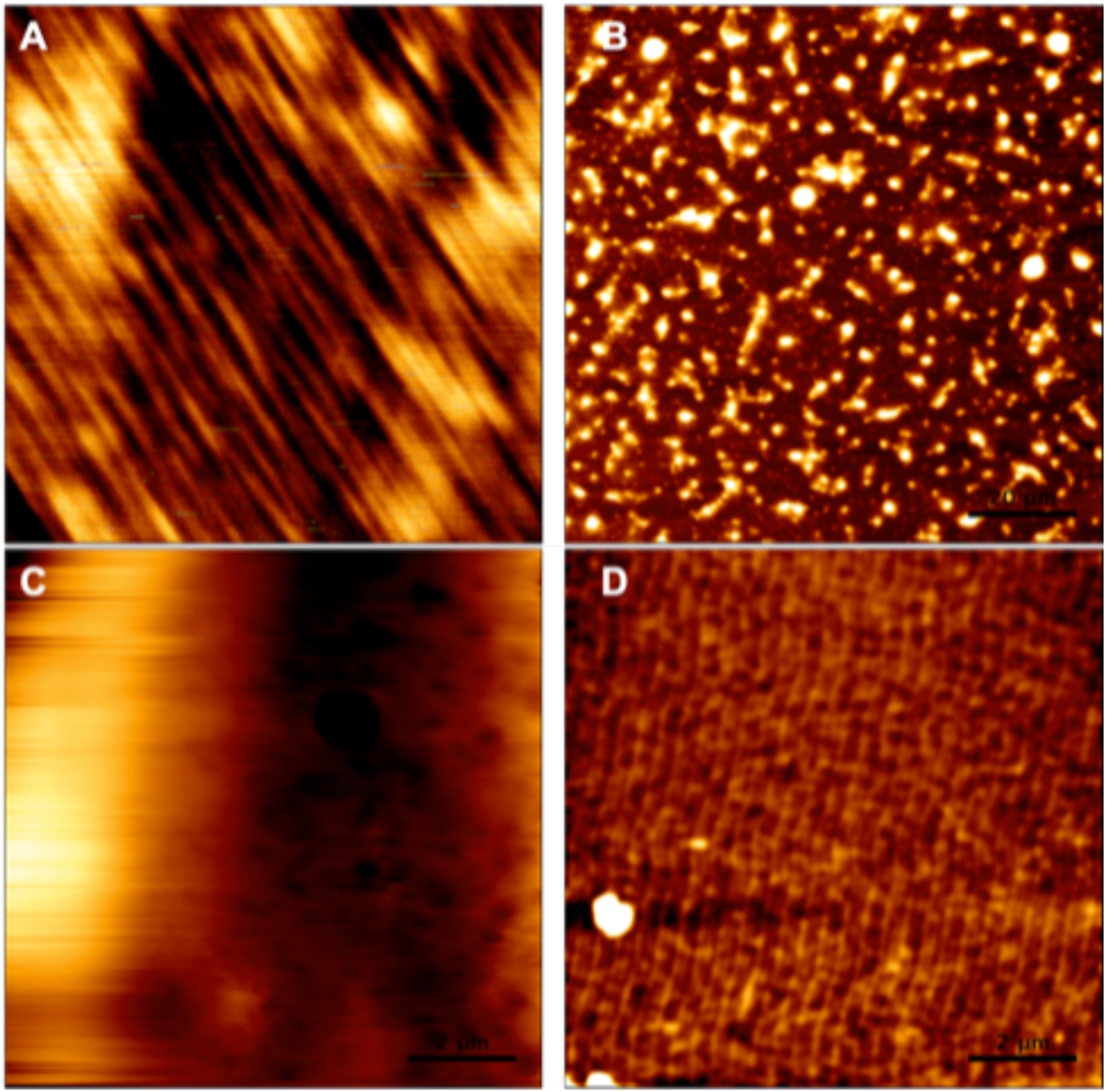
Atomic Force Microscopy (AFM). Representative AFM images. Precipitation of exosomes plus Aβ42 (Figure 3A) shows the formation of long fibers around 20nm tall. The remaining supernatant shows individual structures but no fibers present (Figure 3B). Precipitation of exosomes without Aβ42 present shows soft structures but no fibers (Figure 3C). Precipitation of Aβ42 with no exosomes present (Figure 3D) resulted in empty mica surfaces with only occasional small structures, but no fibers.

Overall these data suggest that exosomes are capable of binding to Aβ peptides in a direct and specific manner. In addition, given the findings of Figure 3, and those of the SIMOA analysis (Figure 1) it appears that Aβ42F has a slower attachment to the exosomes and/or appears to bind at lesser concentrations than the monomeric form.

## Discussion

The ability of exosomes to bind to Aβ40 and Aβ42 amyloid in a selective manner is potentially important for many reasons. APP and Aβ have been shown to accumulate in the MVB [14–16] which is a milieu in which ILV’s that will become exosomes can interact. In addition, exosomes have been reported to be associated with Aβ and subsequently be located in microglia, hence serving as a potential delivery vehicle for Aβ removal [17]. In contrast, the formation of Aβ laden exosomes versus delivering the Aβ cargo to lysosomes for degradation, may serve to seed pathology, a phenomemon observed when the lysosome is dysfunctional [18] allowing the spread of pathology by the Aβ laden exosomes [19–21].

Given these findings, Amyloid/Aβ bound to exosomes can be considered an important plasma measure as a potential biomarker of disease and/or as a read-out for changes in response to therapeutic interventions.

## Acknowledgments

The authors would like to thank Kirk Hansen for his interest in these findings, his suggestions for their interpretation and thoughts on how to proceed with this work. Additionally, the authors would like to thank Tom Anchordoquy for thoughtful feedback and insights on the work presented.

